# Why probe registration is crucial for infant fNIRS and HD-DOT? An evaluation across 49 newborns

**DOI:** 10.64898/2026.10.27.760957

**Authors:** Guy A. Perkins, Ana Fló, Emma Visibelli, Esma Nur Doğanbaş, Natalia Reoyo-Serrano, Eugenio Baraldi, Silvia Benavides-Varela

## Abstract

**Introduction:** Source-detector probe registration, i.e., the mapping of optode positions on the scalp to their true anatomical locations in head/brain space, is crucial for brain imaging with fNIRS or HD-DOT. While individual, MRI-based registration is desirable, it is not always feasible, motivating the development of alternative registration approaches.

**Materials and Methods:** This study uses the STORM-Net registration protocol and quantifies the variation of cap alignment in a cohort of 49 newborns to evaluate how this affects reconstructed images using HD-DOT.

**Results:** Relative to a centrally aligned template, the cohort showed a median probe-position error of 10.12 mm, while the least and most misaligned subjects exhibited mean errors of 4.41 mm and 20.60 mm, respectively.

**Discussion:** The results demonstrate that the level of cap misalignment typically observed in newborn imaging can affect the spatial accuracy of reconstructed functional images, particularly in cases with larger probe displacements. These findings provide a quantitative basis for adopting more conservative interpretations of group-level functional brain activity and functional connectivity results, and strongly support the routine use of registration methods in infant HD-DOT studies.

## 1 Introduction

### 1.1 What is fNIRS and HD-DOT

Functional Near Infrared Spectroscopy (fNIRS) is a non invasive, non ionizing brain, lab and portable based imaging technique, that can measure functional brain activation on the surface of the cortex.^1^ First contrived in 1977,^2^ fNIRS has evolved to encompass myriad applications ranging from clinical medicine,^3–6^ sports science,^7^ neuroscience,^8^ and psychology.^9^ The technique exploits the relative transparency of biological tissue to near-infrared light (650-950 nm) and the differential absorption spectra of oxygenated (HbO) and deoxygenated (HbR) hemoglobin to infer localized changes in the two aforementioned chromophores associated with neural activity.^10^

High-Density Diffuse Optical Tomography (HD-DOT) represents an advanced implementation of fNIRS principles, using a high density array of sources and detectors to enable three-dimensional tomographic reconstruction of haemodynamic changes on the surface of the cortex.^11^ Unlike conventional fNIRS, which uses channel-based measurements between individual source-detector pairs, HD-DOT leverages overlapping measurements, which yield strong sensitivity on the surface of the cortex and tissue modeling to reconstruct volumetric images with improved spatial resolution and depth discrimination.^12–15^ This enhanced capability makes HD-DOT particularly suited for localizing functional brain activity and investigating functional connectivity.^16^

### 1.2 Why on newborns

fNIRS and HD-DOT are ideal imaging modalities on the newborn and infant population, due to their non-invasive and non-ionizing nature, portability, and tolerance to movement.^17^ Traditional neuroimaging techniques such as functional magnetic resonance imaging (fMRI)^18^ or positron emission tomography (PET)^19^ are less practical or require ecological compromises in the experimental procedure.^20, 21^ In the case of fMRI, subjects are required to be in a noisy scanner, with minimal motion and in the case of PET, emit ionizing radiation, which may harm the newborns or infants. Electroencephalography (EEG),^22^ whilst commonly used for infant studies, has limited spatial resolution and is also more sensitive to motion compared to fNIRS. While sedation can address movement, it introduces potential negative long term effects on neurodevelopment outcomes, which limits its suitability for non-clinical research studies.^23^

Nevertheless, neuroimaging research in these developmental populations remains essential, as the neonatal period represents a critical window for brain development, during which maturation of cortical tissue, and the establishment of functional connectivity patterns occur.^24, 25^ Understanding the neural mechanisms underlying early cognitive processes such as language acquisition,^26–30^ memory formation,^31–35^ and social sensory processing^36, 37^ requires neuroimaging techniques that can be safely and repeatedly applied to newborns.^38^ In addition, fNIRS and HD-DOT offer even more effective imaging solutions on these populations due to the thinner scalp and skull thicknesses compared to adults.^39^

### 1.3 Importance and sources of cap misalignment

Accurate spatial localization of the fNIRS derived haemodynamic response function (HRF) and HD-DOT derived volumetric changes in haemodynamics from the cortical surface requires knowledge of the source-detector optode positions relative to underlying brain anatomy. This is because, in fNIRS, the estimated location of brain activity is inferred from the assumed position of each measurement channel. Similarly, in HD-DOT, reconstructed images of brain activity rely on mathematical models (i.e., spatial profile of the sensitive matrix or jacobian) that are highly dependent on the precise placement of the probes on the scalp. As a result, inaccuracies in probe positioning can lead to errors in the estimated location and extent of brain activity.

Typically, probe positions are either assumed from a standard template, which can be provided by the device manufacturer or derived from anatomical head models, or individually estimated using a procedure known as probe registration (or optode localization). Knowing the ‘true’ locations of the probes, or at least minimizing the error between the assumed probe locations and their true position is important for anatomical interpretation of functional results, group-level statistical analyses, and comparison across studies.^40, 41^

However, relying on template probe locations assumes consistent cap placement across participants, which can be particularly challenging in infant studies. Because cap fitting is often performed under time constraints, the positioning of the imaging cap and therefore the location of the probes on the scalp can vary between subjects. In addition, newborns and infants are prone to movement during data collection, which can further shift the cap, especially in subjects with very small or irregular head shapes. Finally, unlike adult participants, newborns and young infants cannot understand or follow verbal instructions to remain still, making it more difficult to minimize movement-related artifacts during data collection.^42^

### 1.4 Current registration methods

There are several approaches to probe registration that have been developed, each with advantages and limitations.^41^ The simplest method assumes a standardized “template” cap placement based on cranial landmarks such as the nasion, inion, the ‘Cz’ vertex of the head and pre-auricular points, following conventions adapted from the 10-20 or 10-5 EEG electrode placement systems.^40, 43^ Then, either through a form of photogrammetry or otherwise, the locations of these anatomical landmarks with the cap on are measured and an affine transformation matrix is calculated using these two sets of landmarks that connect the subject space to the template space^44^. These are computationally straightforward and require no additional equipment, but this approach does not account for individual variations in head size or shape other than where the landmark locations are, which can potentially introduce errors in spatial localization.

Digitization techniques using electromagnetic tracking systems (e.g., Polhemus^4, 45^) capture individual optode positions with on order of millimeter accuracy.^40^ However, the digitizing system, can be time-consuming, and may be challenging to implement with uncooperative infants.^42^ In addition, electromagnetic digitizers may be sensitive to environmental metal interference. The method used for this work is the STORM-Net (Simple and Timely Optode Registration Method) protocol,^46^ which cab be used with infant populations. This method uses a short video of the participant’s head, recorded from the face to the top of the scalp, along with a pre-existing 3D head and cap model to estimate the 3D positions of the optodes. A convolutional neural network identifies anatomical and cap landmarks across the video frames and aligns them with the predefined head model to determine the cap position on the subject’s head. The approach requires only a brief video recording (typically 10 seconds or less), and processing takes only a few minutes per subject. Because the procedure is semi-automated and can be performed using a smartphone camera, it reduces operator-dependent variability and is particularly suitable for newborn and infant studies.

Despite these methodological advances, the extent to which probe misalignment affects functional imaging results in neonatal fNIRS and HD-DOT remains uncharacterized. There has been a study on 4-7 month old infants in which they performed registration between an fNIRS channels to subject specific MRI models in order to evaluate which regions of interest (ROI’s) have reliable registration, however this study did not evaluate the effect of registration vs no registration.^42^ While previous studies have demonstrated the importance of registration in adult populations,^41, 47, 48^ the impact may be amplified in newborns due to their smaller head size, meaning that a given absolute misalignment represents a proportionally larger displacement relative to cortical anatomy.^42^

### 1.5 Aims and objectives

Given the unexplored implications of cap misalignment on neonatal neuroimaging using fNIRS and HD-DOT, the aims and objectives of this study are twofold. The first is to evaluate the range of cap misalignment on a cohort of newborn subjects involved in a typical fNIRS/HD-DOT study, and this will be evaluated by reconstructing registered probe positions using the STORM-net protocol. This misalignment will be calculated by how much the registered probe positions differ from a centrally aligned template. The second is to evaluate how these misalignments affect the imaging of HD-DOT in this cohort of newborn subjects, with particular focus on the best (most similar to the centrally aligned template), worst (least similar to the centrally aligned template) and the median (the middle similarity to the centrally aligned template and most representative of the cohort) case newborn subjects. The reason why these three subject cases were chosen, is so that the effects of misalignment can be reported and discussed across a range of misalignment. By comparing HD-DOT using template probe co-ordinates and registered probe co-ordinates, the ‘real world’ implications of not using subject registration for HD-DOT on newborns can be evaluated.

## 2 Methods

### 2.1 Subject recruitment

Forty-nine healthy full-term newborns were tested in the hospital a few hours to days after birth as part of a larger research project aimed at investigating early cognitive functions. The demographic and clinical information of the whole sample, as well as the three case study subjects used for this work can be seen in table 1. All participants were in absence of cephalohematoma or other conditions that could affect cortical hemodynamics, and they all had intact hearing. Newborns were recruited from the neonatal care of obstetric division of the University Hospital of Padova, Italy. Informed consent was obtained from all parents, and the protocol was approved by the Ethics Committee for Clinical Research of the Province of Padova (Italy) They were held by their mothers or were lying in their cribs, which is denoted in the bottom row of table 1.

**Table 1.** A summary of the demographic and clinical information, for the full sample, the best case subject, median case subject and worst case subject in terms of probe alignment respectively. Gestational age (GA) given by the number of weeks (W) plus the number of days (D). BW = Birth weight in grams. Head Circ. = Head Circumference in centimeters. Values presented are in the format median (IQR 1-3), where appropriate.

| Variable | All Subjects | Best case | Median case | Worst case |
|---|---|---|---|---|
| N Subs. | 49 | 1 | 1 | 1 |
| Sex (M:F) | 22:27 | 0:1 | 0:1 | 0:1 |
| GA (W+D) | 39+1 (38-40) | 37+3 | 38+3 | 40+5 |
| BW (g) | 3150 (2870-3425) | 3150 | 2320 | 2970 |
| Head Circ. (cm) | 34.0 (33.5-35.25) | 34 | 31.5 | 32.7 |
| Age at testing (days) | 2 (1-2) | 6 | 2 | 3 |

### 2.2 Data Acquisition and Probe Placement

Data were acquired using LUMO, a modular high-density diffuse optical tomography (HD-DOT) system developed by Gowerlabs Ltd. (UK). The system operates at a frame rate of 10 Hz and it is composed of multiple independent sensor modules, or tiles, which together form a dense network of light sources and detectors.

Each hexagonal tile contains three dual-wavelength LED sources (735 and 850 nm) and four photodiode detectors. Measurements within a tile are obtained at source–detector separations of approximately 10 mm and 20 mm, while measurements between tiles are acquired throughout the whole array. Each tile measures 29 mm across at the widest point.

For this study, a layout consisting of 15 tiles was used, which yielded a total of 2700 logical channels per wavelength. Of these, approximately 556 were expected to achieve source-detector separation with acceptable signal quality (*≤* 40 mm separation). Six tiles were positioned in the fronto-temporal and partial areas in each hemisphere, and 3 positioned in the pre-frontal cortex. A black stretchy cap was used to attach the tiles. The optode arrangement and the corresponding channels are shown in figure 1.

**Fig 1.**
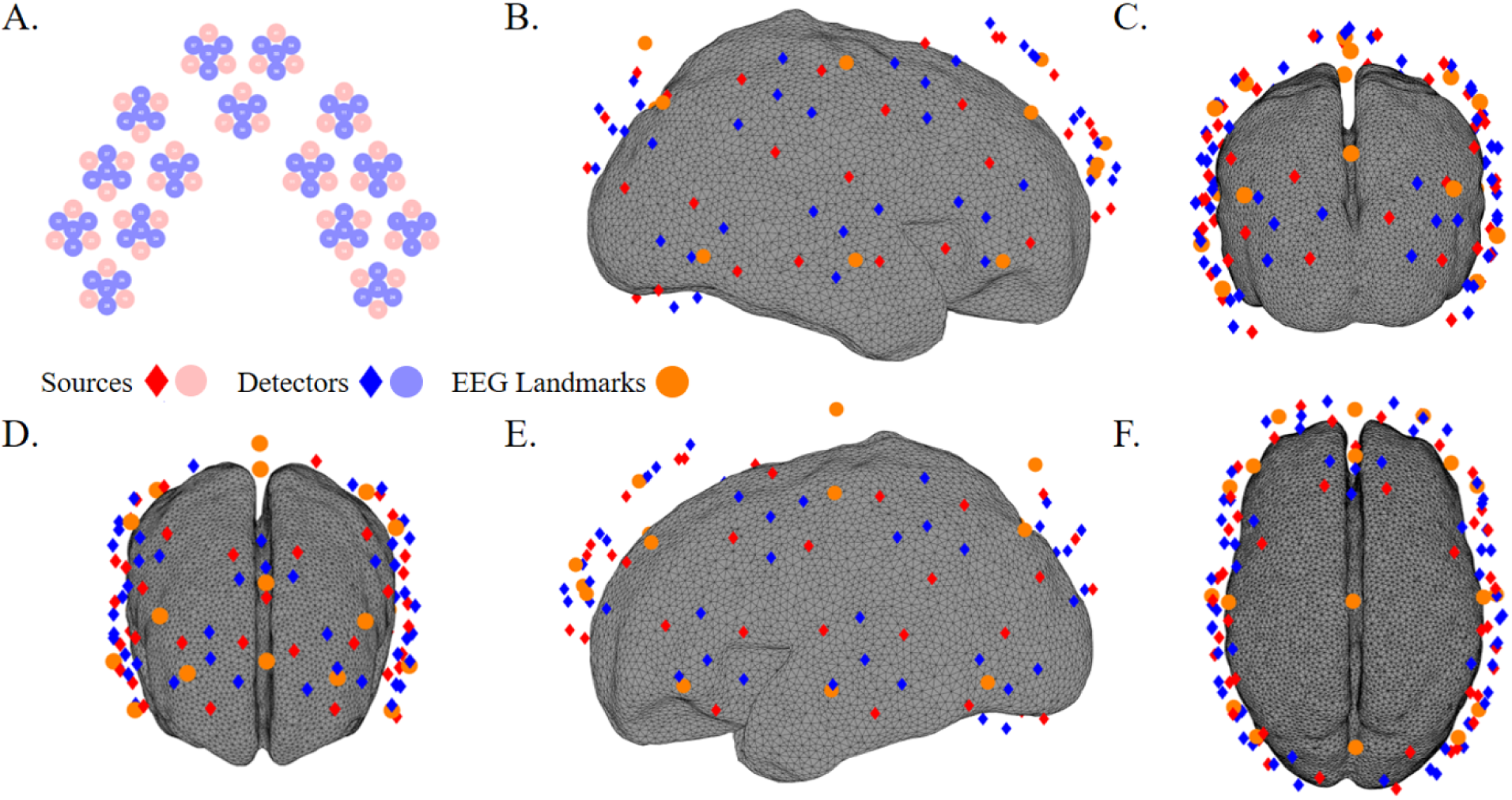
The source-detector probe set up for the experiment. A. The 45 sources and 60 detectors shown in 2D. These were derived from the raw .lufr file from the LUMO system. B-F. Newborn smoothed gray matter surface model with the source-detector probes and selected EEG (10-5) landmarks shown for reference. These are AFz, AF3, AF4, F7, F3, Fz, F4, F8, T7, C3, Cz, C4, T8, P3, Pz, P4, P7 and P8. B. The right hemisphere. C. Posterior of the cortex. D. Anterior of the cortex. E. The left hemisphere. F. Top view of the cortex.

### 2.3 Cap positioning and Registration technique

Before starting the experiment, newborns’ head circumference were measured using a flexible nonstretch measuring tape placed around the widest part of the head, passing above the eyebrows and ears and around the occipital prominence.

After the measurements were obtained, the cap was positioned on the newborn’s head using the bilateral preauricular points as anatomical references to ensure consistent alignment with the anterior edge of the central tile. Moreover, the bottom row of tiles was aligned approximately along the F7–T3–T5 and F8–T4–T6 trajectories in the left and right hemispheres respectively. The anterior tiles were placed over Fp1 and Fp2, ensuring that the cap was symmetrically positioned relative to the face and that the front edge followed the curve of the eyebrows.

The STORM-NET tool^46^ was used to digitize the positions of the optodes. STORM-NET determines the locations of optodes and head markers from a short video recorded from the front of the head, following a 90-degree arc to the vertex. This approach is particularly suitable for newborns, as the video can be rapidly recorded while the infant lies in their crib. The effectiveness of STORM-NET relies on a series of offline steps that must be performed before data acquisition. These include creating a 3D head model (figure 2A), generating synthetic data that replicates the geometry and movement of a head wearing the cap, and training a neural network to estimate sensor positions from video data (figure 2B). Once trained, the model can estimate sensor positions using 10 frames extracted from short videos.

**Fig 2.**
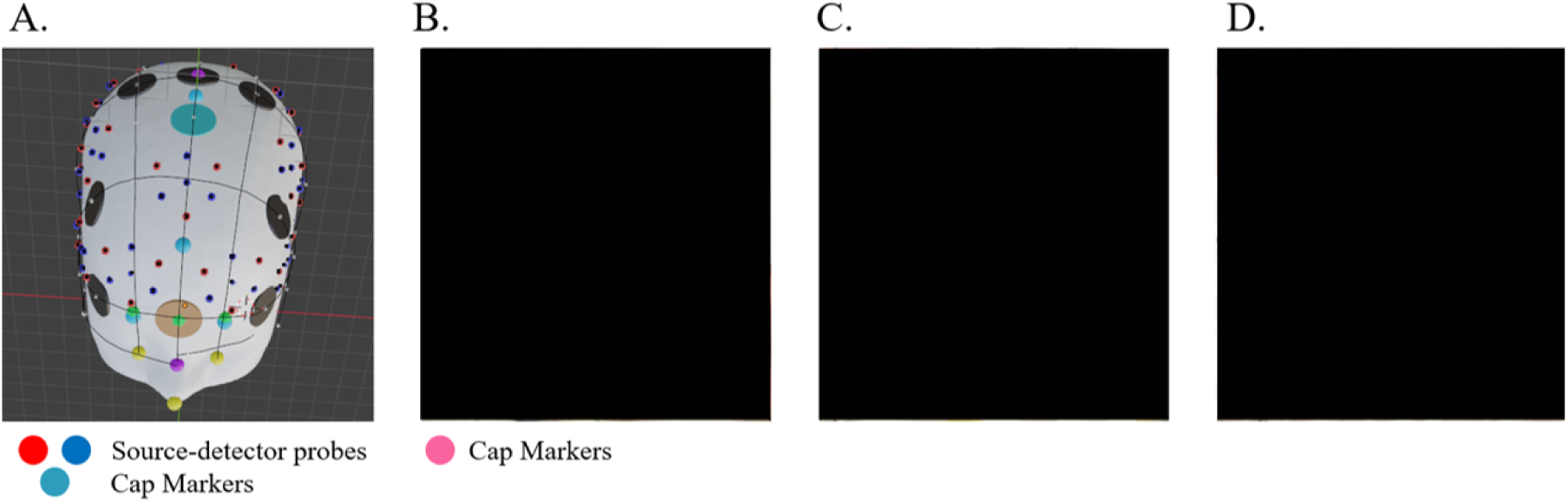
A. The 3D model created and used as part of the Storm-net registration process. B-D. Three frames from the 10 second video required for the Storm-net registration method. These have been redacted as per the requirements for this pre-print.

The 3D head model has to be generated for each cap and needs to have 4 cap markers (left triangle, middle triangle, right triangle, and top), 3 head markers (left eye, right eye, and nose tip), at least four standard locations (Cz, Fp1, Fp2, Fpz), and the locations of the probes to be digitised. The original STORM-NET paper proposes using a digitising device to capture positions with the cap placed on a head mannequin. Although digitalisation of HD-DOT probe positions is possible^49,16^ it is complex due to the high number of probes. In this study, a digital model was created directly, capitalising on the fact that each LUMO cap is designed with fixed tiles and their positions are already digitalised for registration. Indeed, each LUMO recording already contains theoretical positions for all optodes and head markers (nasion, vertex, right and left auricular). Additionally, Gowerlabs provided the head render matching the head size used in the cap design. To create the digital head model, the head render, optode positions, and head marker positions were loaded into Blender 3.6.5 (Amsterdam, Netherlands). The head was aligned with the head markers, and the positions of the International 10-20 system were identified by drawing the midline between nasion and inion, preauricular to preauricular line, skull circumference, and the required lines at specified intervals. Thus, procedures usually performed on a subject or a mannequin were performed on a digital mannequin. The four cap markers required by STORM-NET were placed in both the digital model and the physical cap (Teal markers in Figure 2A and pink markers in figure 2B). The left triangle marker was placed on Fp1, the right triangle marker on Fp2, and the middle triangle on Fz, while the top marker was placed between Cz and FCz due to space limitations in tiles and electrode placement. The positions of the optodes, standard locations, cap markers, and head markers were then extracted and saved in a text file, which served as input to generate the STORM-NET model.

During each recording session, videos were recorded using a phone in the horizontal position, following a path covering 90 degrees of the newborn’s head from the front of the face to the vertex. The cap markers, as well as the eyes and nose tip, were manually identified in the 10 frames selected by STORM-NET to estimate the actual optode placement.

### 2.4 Paradigm

Neonates were tested in a dimly lit hospital room while lying in their cribs, in quiet rest or sleeping, to ensure their comfort and maintain an ecologically valid environment. They completed a modified version of previous experimental designs to assess memory for words in neonates^33^.^32^ The paradigm consisted of 4 phases: 2 phases of auditory stimulation and 2 phases of silence that were intermixed (i.e, silence-stimulation-silence-stimulation). Each phase lasted 5 minutes for a 20 min test session in total. Sound stimuli were presented in blocks lasting around 10s and separated by 25-35s. The stimuli was delivered using the PsychoPy software^50^ via two loudspeakers located at a distance of 70cm from the newborns’ head. A video camera was used to monitor the behavior of the newborns and the placement of the probes throughout the experiment.

### 2.5 Pre-processing and Image reconstruction

The pre-processing pipeline consisted in converting raw intensity to optical density (OD), conversion and quality assessment of individual channels based on signal-to-noise ratio (SNR) (SNR *≥* 12) and scalp coupling index^51^ (SCI) (SCI *≥* 0), and then motion artifact detection using a modified Homer3^52^ function called ‘ot MotionDetection hmr3’ (see shared code) with variables thresh = 2.0 and tMotion = 2.0, and correction using spline correction^53^ (p=0.99) and wavelet correction^54^ (iqr parameter = 1.5) as implemented in Homer3. Then bandpass filtering (0.01–0.5 Hz) was used to remove slow drift and high-frequency noise,^55^ short-channel regression (SDS *≤* 10 mm) was used to remove systemic physiological confounding from all channels^56^ and then the data was down-sampled at 1Hz. Following pre-processing, the processed OD data were organized into blocks corresponding to blocks of the auditory stimuli. These OD data were then block averaged using a baseline of −5s to −1s prior to the start of the stimuli and the 20s post stimuli onset.

The HD-DOT image reconstruction composed of two parts, the first was the jacobian (or sensitivty, or “A”) matrix generation, i.e modeling forward propagation of photon diffusion in tissue and the second was solving the inverse problem for the actual 3D image reconstruction. These were both performed using NIRFAST (Near Infrared Fluorescence and Spectral Tomography), an open-source finite element method software package for modeling light transport in tissue.^57^

The jacobians were generated using the 4D neonatal head finite element models^58^ (FEM’s) which are a set of head models spanning 29 to 44 weeks GA, containing a model for each unique week and were created from the average of 324 T2 MRI images. Jacobians and image reconstruction were performed for all participants, but are displayed here only for the best, median and worst case subjects which used the 37, 38 and 41 week GA model’s respectively. The optical properties used for these models can be seen in table 2, and a refractive index of n = 1.4 was used.

**Table 2.** The optical properties used for the different tissues in the mesh Finite Element Model (FEM).^16^ CSF = cerebral spinal fluid, GM = gray matter, WM = white matter. All values given in units of mm*^−1^*.

| Layer | *μ*<sub>*a*</sub> 850 nm | *μ*<sub>*a*</sub> 735 nm | *μ*′<sub>*s*</sub> 850 nm | *μ*′<sub>*s*</sub> 735 nm |
|---|---|---|---|---|
| Scalp | 0.020 | 0.015 | 0.751 | 0.876 |
| CSF | 0.004 | 0.002 | 0.300 | 0.300 |
| GM | 0.019 | 0.018 | 0.673 | 0.860 |
| WM | 0.021 | 0.016 | 1.011 | 1.218 |

Then jacobians were created for each of the three test case subjects, with two jacobians created for each subject, the first using the template centrally aligned source-detector positions and the second using the registered probe positions from the STORM-net protocol. Then, the inverse problem was solved using NIRFAST’s regularized least-squares approach^57^ with Tikhonov regularization (*λ* = 0.01) to produce 3D volumetric images of haemoglobin concentration changes in oxygenated (HbO) and deoxgynated (HbR) hemoglobin respectively.^59^ This means for each of the test case subjects, there were two sets of volumetric changes in HbO (ΔHbO) and HbR (ΔHbR), one from the template probe jacobian and one from the registered probe jacobian.

### 2.6 Error calculation of the misalignment

For each subject, the 3D optode positions obtained from STORM-Net registration were compared to the corresponding template positions in order to quantify the difference between the two sets of probe positions. These misalignment ‘errors’ were computed as the Euclidean distance between each registered optode position and the corresponding template probe position, given by equation 1,

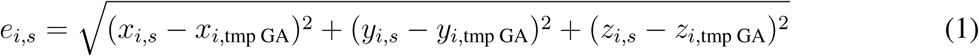

where *e_i,s_*is the misalignment error for optode *i* and subject *s*, and *x_i,s_, y_i,s_, z_i,s_* are the registered coordinates and *x_i,tmp GA_*, *y_i,tmp GA_*, *z_i,tmp GA_* are the template coordinates corresponding to the GA of each given subject.

To quantify the spatial differences from functional activation, ΔHbO was considered at t=8s after the stimulus onset, this time point was chosen based upon the features of the newborn HRF.^60^ Clusters of ΔHbO were calculated, with their 3D centroid locations recorded. This was done by creating masks of positive and negative ΔHbO, in which nodes were included in the mask if they had absolute values of ΔHbO *>*50% of the maximum change in ΔHbO. Clustering was then performed using the ‘graph’ and ‘conncomp’ functions in matlab with these masks and ΔHbO to create clusters for positive and negative changes and the template and registered derived ΔHbO. The displacement between clusters was calculated by the euclidean distance between their centroids and the dice coefficient was calculated to quantify the spatial overlap between clusters and is given by equation 2,

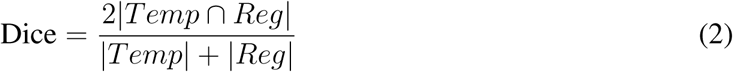

where *Temp* and *Reg* represent the nodes of corresponding clusters from template and registered ΔHbO respectively, such that the Dice coefficient ranges from 0 (no overlap) to 1 (perfect overlap). Cluster comparisons were excluded from analysis if the clusters had Euclidean distances between centroids of *>*50 mm or ratios of the maximum absolute value of ΔHbO *<* 0.5 or *>*2, in order to prevent comparisons between clusters which had erroneously been classified, since these parameters represent realistic changes between clusters. The probe misalignment and changes in the clusters of ΔHbO were then visualized using the 4D infant head model for each of the three test case subjects and are presented in the results section.

### 2.7 Probe displacement correlations

Finally, in order to investigate if external factors contribute to the probe alignment error, linear correlations were computed between the mean alignment error and several different metadata from the full sample of participants. These were the head circumference (cm), the age of the newborn when the HD-DOT data was acquired (days), their birth weight (gr), their gestational age (days) and then two types of data from the LUMO motion sensors. Each tile on the LUMO device has a motion sensor which measures the angular velocity (units of deg s*^−^*^1^) and the angular acceleration (units of g, the acceleration due to gravity (ms*^−^*^2^)). Both of these values are given in the x, y and z orthogonal axis, therefore a magnitude vector was computed for these values by taking the euclidean norm, averaged across all tiles and averaged across time for each subject, to give *|ω|*, the magnitude of the angular velocity vector and *|α|*, the magnitude of the angular acceleration vector. These metadata were selected based upon their plausibility of effecting initial cap placement, due to head size or movement, and the cap moving during the experiment due to movement.

## 3 Results

### 3.1 Probe displacement

Figure 3 shows the distributions of the displacement between the template and registered probes (for each given source and detector) for the best, median and worst case scenario. Each subject has 105 probes and there are in total 5145 probes for all 49 subjects, with each data point representing a source or detector. These three highlighted subjects were chosen by ordering the average probe displacement of all 49 subjects, with the best (lowest average displacement), median (median average displacement) and worst (highest average displacement) case subjects. A paired t-test between the best, median and worst case probe displacements showed that there were statistically significant differences between the best, median and worst case distributions respectively.

**Fig 3.**
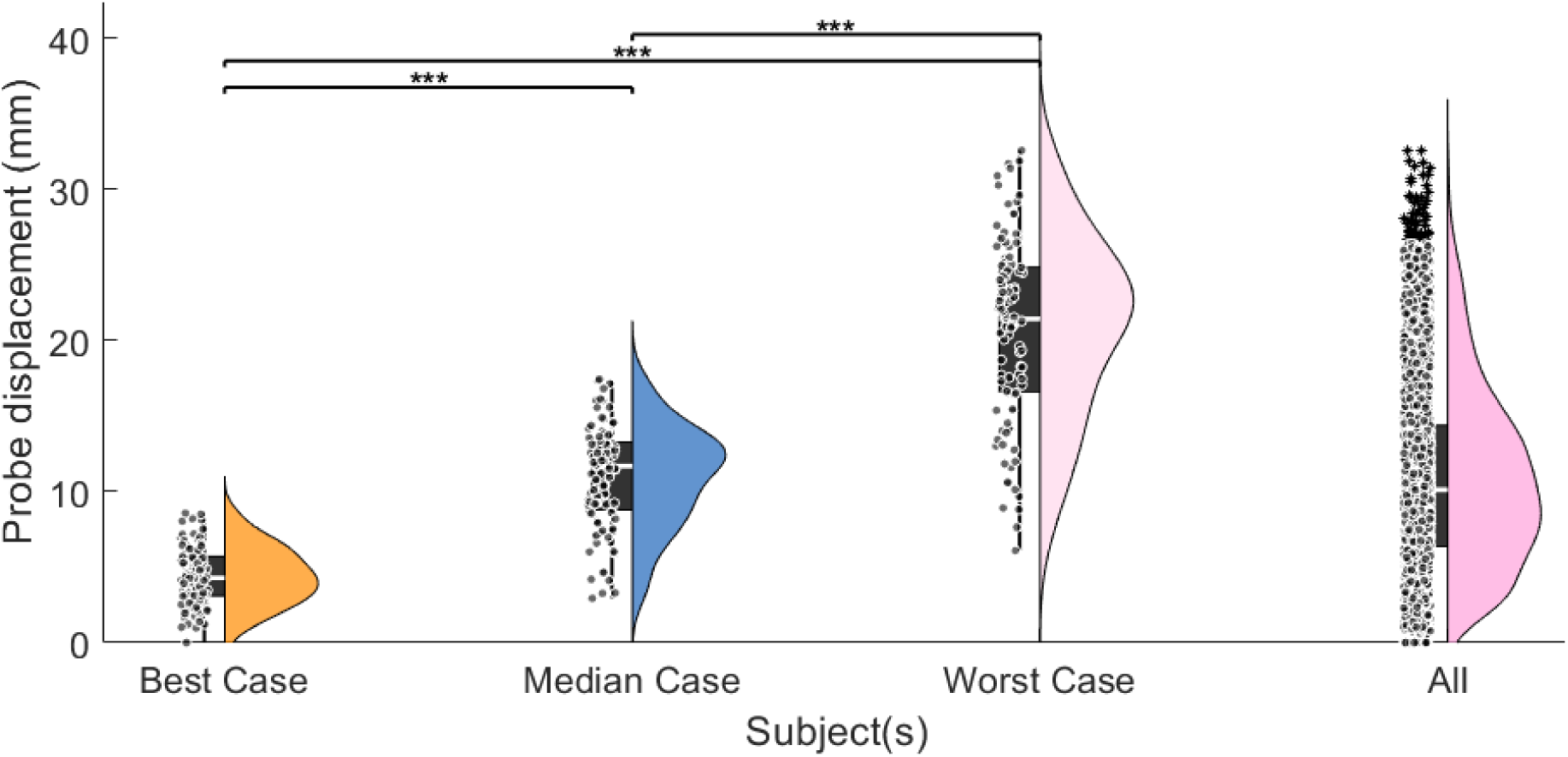
Distributions of the displacement between the template and registered probes (for each given source and detector) for the best, median and worst case scenario subjects respectively, as well as for all 49 subjects. *** denotes a p value of *<* 0.001 from a paired t-test, after corrections for multiple comparisons.

Figure 4 shows the template probes and registered probes for the best case subject (median probe displacement = 4.29 mm). Generally the cap was well aligned, which can be seen in the image of the newborn, figure 4A. The cyan line runs from the nose of the newborn, then goes through the nasion and between the centre of their eyes. This axis is meant to depict a visual guide of what an anatomical central alignment would look like on this subject, which is what a centrally aligned probe template assumes and is therefore called the ‘template axis’. The other line, coloured magenta, starts from the same point as the template axis, however it goes through the centre reference point on the cap (green dot) which is one of seven reference points used for the STORM-NET registration process. This axis is meant to visually depict the true alignment of the cap, and therefore where the registered source-detector probes are, with respect to the centrally aligned template axis, and therefore is called the ‘registered axis’.

Figure 4B. shows that the displacement between the template and registered probes are very small in the three frontal tiles (less than 2 mm). The displacements increase on both the left and right hemisphere in the central and temporal areas (2-6 mm; Figures 4C and D). The largest displacements can be observed in the peripheral tiles, with the largest displacements (8 mm) occurring on the right hemisphere.

The median case subject can be seen in figure 5, and as with the best case subject, the registered axis is displaced to the right (left in image) of the template axis. In addition, the cap further back along the newborn head than what the centrally aligned template assumes (Fig. 5B to D). Overall, the median displacement for this case subject is 11.68 mm with smaller displacements for probes on the left hemisphere of between 4 to 12 mm, figure 5C, compared to 12 to 14 mm on the right hemisphere, figure 5D.

**Fig 4.**
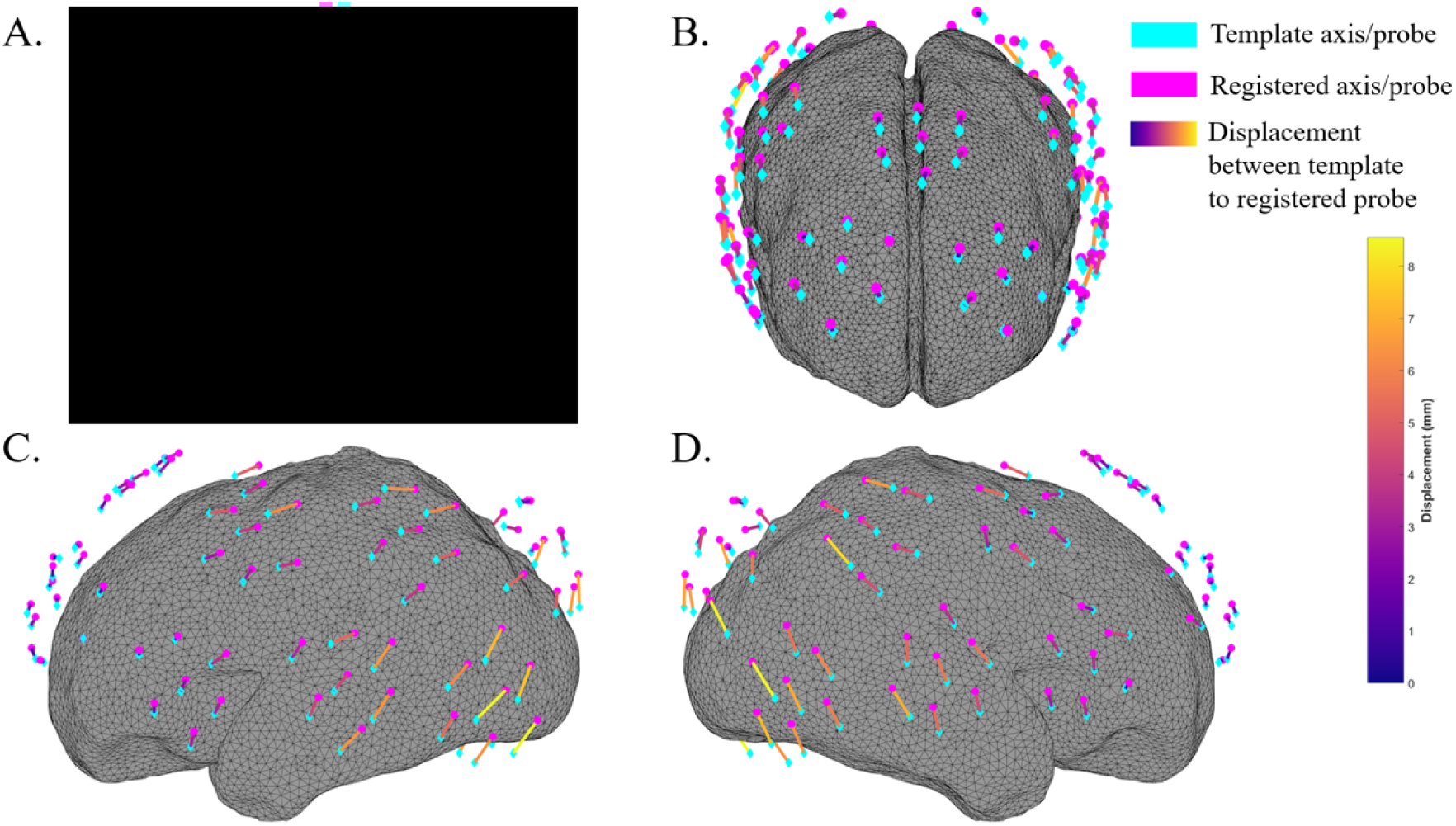
The best case subject. A. An image of the best case subject showing the cap alignment, along with lines depicting the anatomically centrally aligned ‘template’ axis from the center of the nose through the central green dot marker and a ‘registered’ axis from the center of the nose through the center of the central frontal tile on the lumo cap. These have been redacted as per the requirements for this pre-print. B-D. The source-detector probes of from the template and from after registration, along with lines showing the displacement between each corresponding source-source and detector-detector, displayed on the surface of the scalp (hidden) above a smoothed gray matter surface head model^58^ (37 week GA).

**Fig 5.**
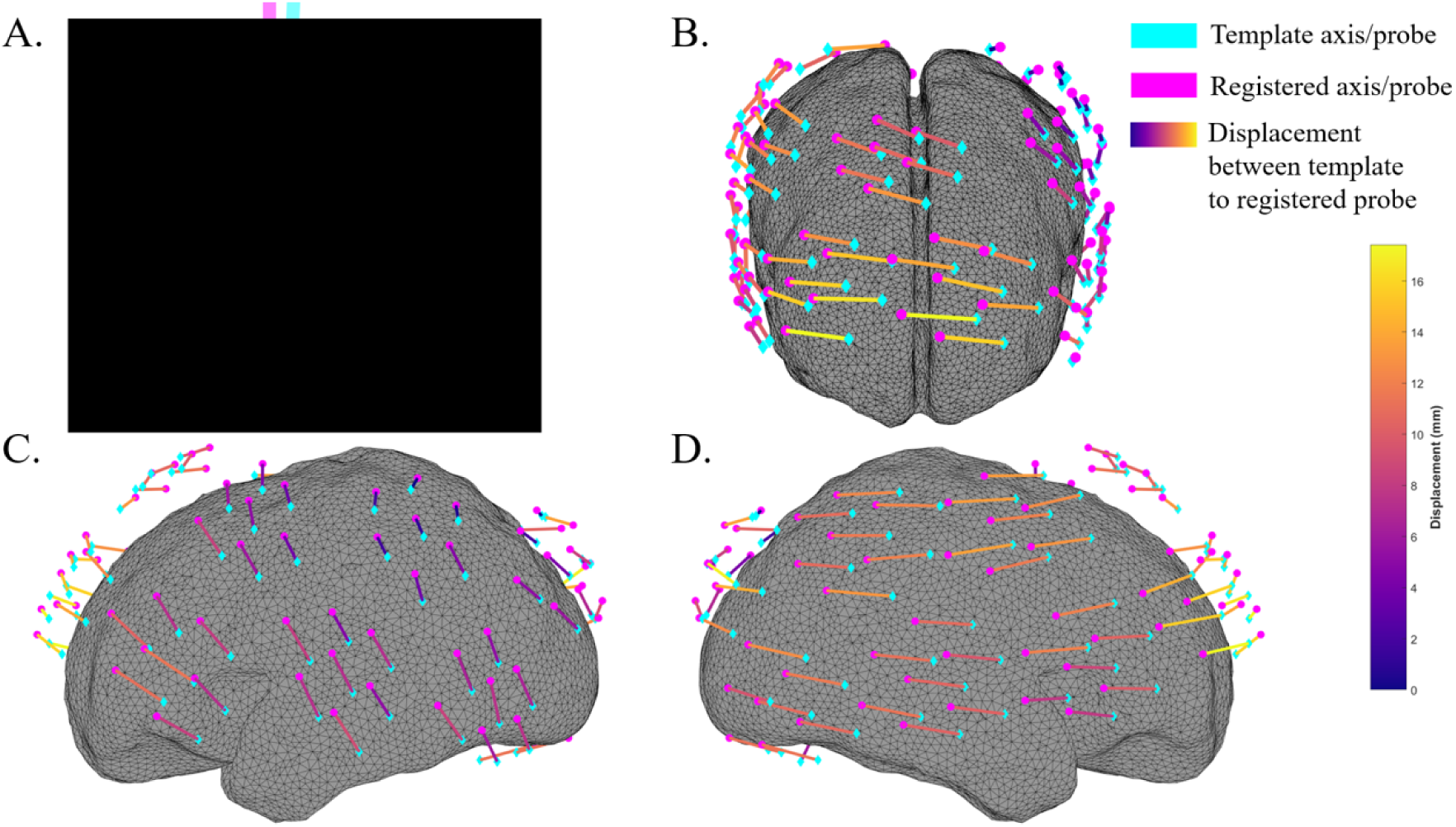
The median case subject. A. An image of the best case subject showing the cap alignment, along with lines depicting the anatomically centrally aligned ‘template’ axis from the center of the nose through the central green dot marker and a ‘registered’ axis from the center of the nose through the center of the central frontal tile on the lumo cap. These have been redacted as per the requirements for this pre-print. B-D. The source-detector probes of from the template and from after registration, along with lines showing the displacement between each corresponding source-source and detector-detector, displayed on the surface of the scalp (hidden) above a smoothed gray matter surface head model^58^ (38 week GA).

It can also be observed that the orientation of the displacement differs between areas of the scalp. On the left hemisphere, the displacement across tiles appears to be nearly vertical as viewed in figure 5C, and the frontal 3 tiles share this orientation as can be seen in figure 5A, whereas orientation of displacement appears more horizontally aligned on the right hemisphere, as viewed in figure 5D.

Finally, the worst case subject can be seen in figure 6. As with the median case, the largest displacement of probes are observed in the 3 frontal tiles (25-32 mm; figure 6B). The direction of displacements are consistent between the hemispheres and frontal area, and unlike the median case, here there are larger displacements on the left hemisphere, figure 6C, of 20 to 25 mm, compared to 5 to 15 mm displacements on the right hemisphere, figure 6D. Overall, the median displacement on the worst case subject is 21.42 mm, which is nearly double that of the median case and a factor of 5 larger than that of the best case subject.

**Fig 6.**
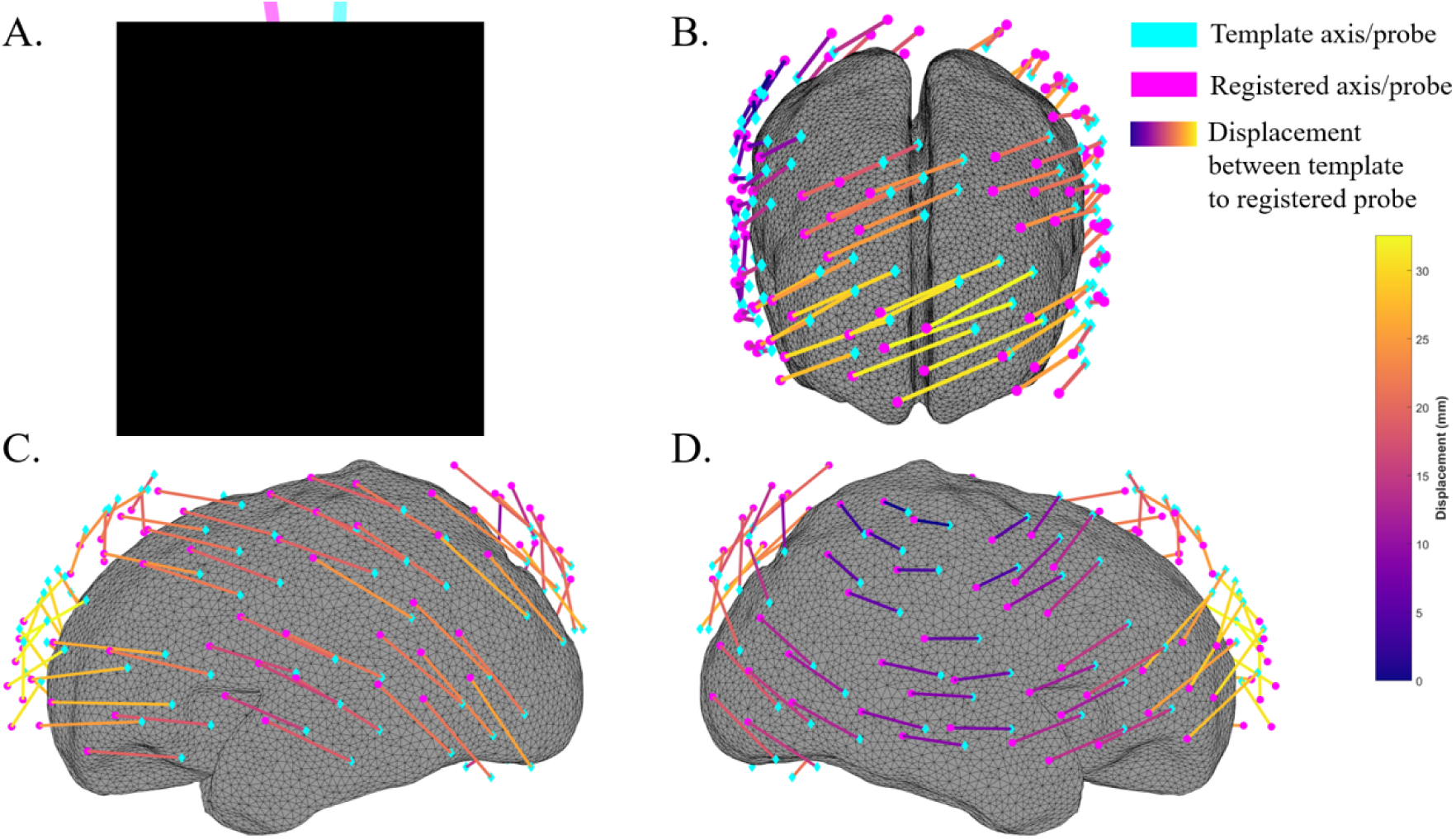
The worst case subject. A. An image of the best case subject showing the cap alignment, along with lines depicting the anatomically centrally aligned ‘template’ axis from the center of the nose through the central green dot marker and a ‘registered’ axis from the center of the nose through the center of the central frontal tile on the lumo cap. These have been redacted as per the requirements for this pre-print. B-D. The source-detector probes of from the template and from after registration, along with lines showing the displacement between each corresponding source-source and detector-detector, displayed on the surface of the scalp (hidden) above a smoothed gray matter surface head model^58^ (41 week GA).

### 3.2 Imaging displacement

The image reconstructions of changes in HbO are shown for the best case subject in figure 7, which shows that the centroid of the clusters of ΔHbO (focal activity of HbO changes) in the template and the registered probes is very similar. The metrics of these clusters are given in table 3.

**Fig 7.**
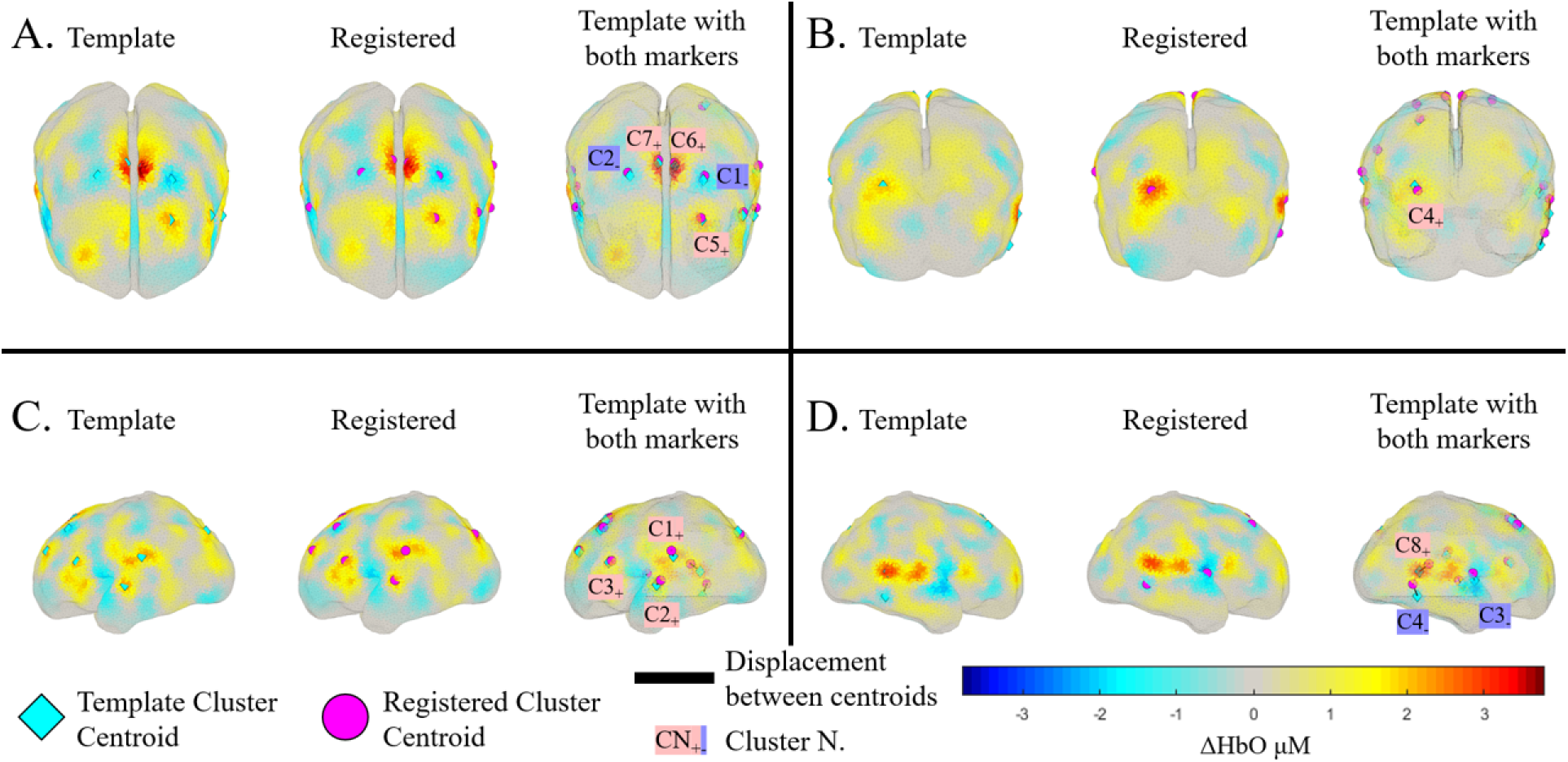
Image reconstruction of ΔHbO from HD-DOT measurements on the best case subject at t = 8 s during a block of auditory stimulation. Each panel A-D shows the location of the significant clusters calculated from using the template probe positions for the Jacobian generation in the first image (highlighted in cyan) and from using the registered probe positions in the second image (highlighted in magenta). The third image in each panel shows the template ΔHbO map overlaid with labels indicating the clusters identified in both the template and registered cases, which are linked with a line for comparative purposes. Labels in pink indicate clusters with significant increments in HbO. Labels in purple indicate clusters with significant decrements in HbO. Each panel shows a different view of the smoothed gray matter surface model, A. Top, B. Posterior, C. Lateral Left and D. Lateral Right. Metrics of these clusters are shown in table 3.

**Table 3.** Metrics for each cluster of ΔHbO from the best case subject in figure 7. These metrics are the displacement between the centroid of the ΔHbO clusters from the template to the registered images (euclidean distance) and the dice coefficient between these.

| Cluster N<sub>+</sub> | Displacement (mm) | Dice | Cluster N<sub>−</sub> | Displacement (mm) | Dice |
|---|---|---|---|---|---|
| 1 | 3.14 | 0.21 | 1 | 2.01 | 0.70 |
| 2 | 3.28 | 0.24 | 2 | 1.73 | 0.44 |
| 3 | 1.63 | 0.53 | 3 | 4.29 | 0.52 |
| 4 | 2.08 | 0.32 | 4 | 6.82 | 0 |
| 5 | 0.92 | 0.83 |  |  |  |
| 6 | 1.59 | 0.85 |  |  |  |
| 7 | 1.68 | 0.87 |  |  |  |
| 8 | 6.31 | 0.32 |  |  |  |

Changes in HbO from the template and registered probe positions for the median case subject can be seen in figure 8, and unlike for the best case subject, the displacements between the centroids of ΔHbO between the template and registered probes are appreciable. This is corroborated by the metrics of these clusters shown in table 4, given that the mean displacement of the positive and negative clusters are 12.07 mm (*σ* = 3.97 mm) and only 1 cluster out of the 8 has a non zero dice coefficient. Of particular note is negative ΔHbO cluster number 2, which for using the template probes has its centroid on the central left hemisphere on the surface of the cortex, whereas for the registered probes, its centroid moves to the central right hemisphere. This is the most visually apparent change of spatial location, however 5 of the 8 clusters for this subject have displacements larger than negative ΔHbO cluster number 2, meaning equal, if not more attention should be paid to other cluster’s discrepancies.

**Fig 8.**
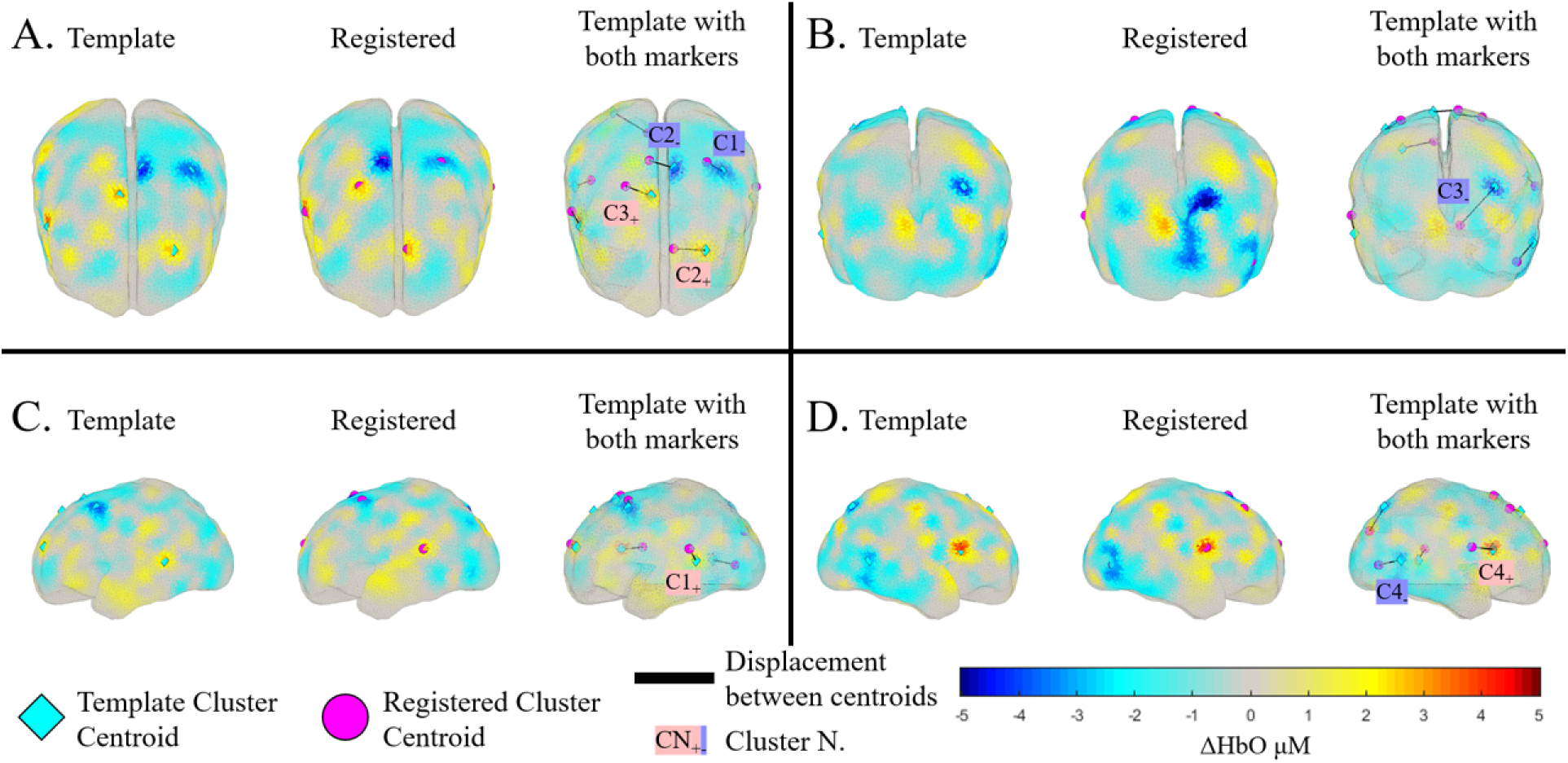
Image reconstruction of ΔHbO from HD-DOT measurements on the median case subject at t = 8 s during a block of auditory stimulation. Each panel A-D shows the location of the significant clusters calculated from using the template probe positions for the Jacobian generation in the first image (highlighted in cyan) and from using the registered probe positions in the second image (highlighted in magenta). The third image in each panel shows the template ΔHbO map overlaid with labels indicating the clusters identified in both the template and registered cases, which are linked with a line for comparative purposes. Labels in pink indicate clusters with significant increments in HbO. Labels in purple indicate clusters with significant decrements in HbO. Each panel shows a different view of the smoothed gray matter surface model, A. Top, B. Posterior, C. Lateral Left and D. Lateral Right. Metrics of these clusters are shown in table 4.

**Table 4.**
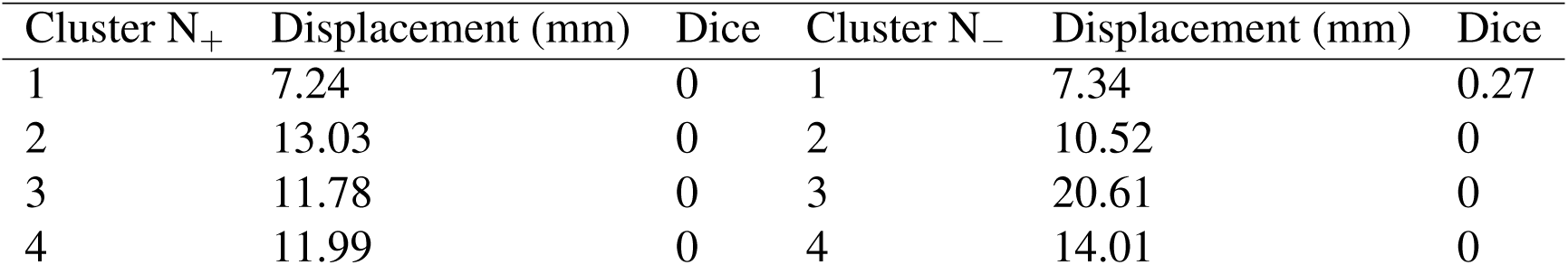
Metrics for each cluster of ΔHbO from the median case subject in figure 8. These metrics are the displacement between the centroid of the ΔHbO clusters from the template to the registered images (euclidean distance) and the dice coefficient between these.

Finally, changes in HbO for the worst case subject can be seen in figure 9 and as with the median case subject, the displacement between the centroids of ΔHbO clusters is visually noticeable, with a mean displacement of 17.94 mm (*σ* = 4.95 mm), with metrics of individual clusters shown in table 5. The clusters are grouped together in four broad locations, across the frontal central cortex (positive clusters 1, 3 and 4). This is largely consistent with the probe displacements seen in figure 6 with the largest displacements seen in and around the frontal three tiles.

**Fig 9.**
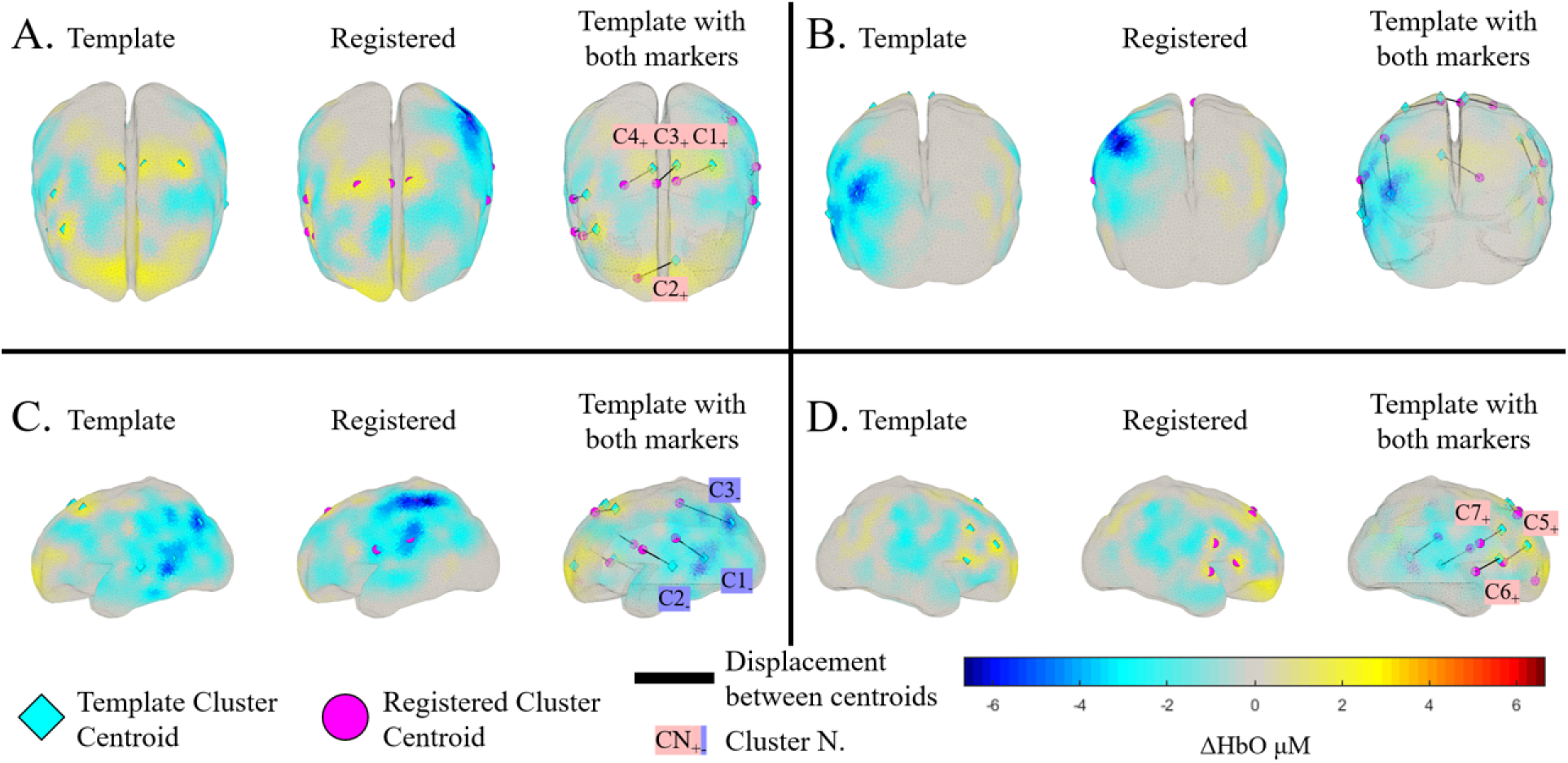
Image reconstruction of ΔHbO from HD-DOT measurements on the worst case subject at t = 8 s during a block of auditory stimulation. Each panel A-D shows the location of the significant clusters calculated from using the template probe positions for the Jacobian generation in the first image (highlighted in cyan) and from using the registered probe positions in the second image (highlighted in magenta). The third image in each panel shows the template ΔHbO map overlaid with labels indicating the clusters identified in both the template and registered cases, which are linked with a line for comparative purposes. Labels in pink indicate clusters with significant increments in HbO. Labels in purple indicate clusters with significant decrements in HbO. Each panel shows a different view of the smoothed gray matter surface model, A. Top, B. Posterior, C. Lateral Left and D. Lateral Right. Metrics of these clusters are shown in table 5.

**Table 5.** Metrics for each cluster of ΔHbO from the best case subject in figure 9. These metrics are the displacement between the centroid of the ΔHbO clusters from the template to the registered images (euclidean distance) and the dice coefficient between these.

| Cluster N<sub>+</sub> | Displacement (mm) | Dice | Cluster N<sub>−</sub> | Displacement (mm) | Dice |
|---|---|---|---|---|---|
| 1 | 18.50 | 0 | 1 | 19.53 | 0 |
| 2 | 18.23 | 0 | 2 | 19.08 | 0 |
| 3 | 11.64 | 0 | 3 | 30.93 | 0 |
| 4 | 13.50 | 0 |  |  |  |
| 5 | 17.03 | 0 |  |  |  |
| 6 | 15.65 | 0 |  |  |  |
| 7 | 15.35 | 0 |  |  |  |

The largest displacement is given by negative ΔHbO cluster 3, 30.93 mm, which is due to a similar phenomena as seen for the median case subject, negative ΔHbO cluster 3 in figure 8, whereby in the template ΔHbO image, the cluster is characterized by a distinct spot of a larger decrease in HbO, figure 9C left peripheral area. Whereas, in the registered case, the cluster merges with nearby decreases in HbO, and therefore the centroid of the cluster is given in the middle of this larger cluster, thus the displacement between centroids increases more than the proportion of relative probe displacement. The displacement is such that the centroid goes from being in the left peripheral area to the left central area of the cortex.

In addition, two clusters, positive clusters 2 and 3 go from having centroids in the left hemisphere for the template probe case, to having centroids of ΔHbO in the right hemisphere, figure 9A. To highlight how large these displacements are relative to the newborn, it should be recalled that the head circumference of this subject was 32.7 cm, and given that the largest displacement of ΔHbO was 30.93 mm, which strikingly is approximately 10% of this subject’s head circumference. Finally, it should be noted that all clusters for the worst case subject have a dice coefficient of 0, which means that there was no spatial overlap of the clusters of ΔHbO.

### 3.3 Probe displacement correlations

Correlations between experimental metadata and probe alignment error are shown in figure 10. None of the computed correlations were statistically significant.

**Fig 10.**
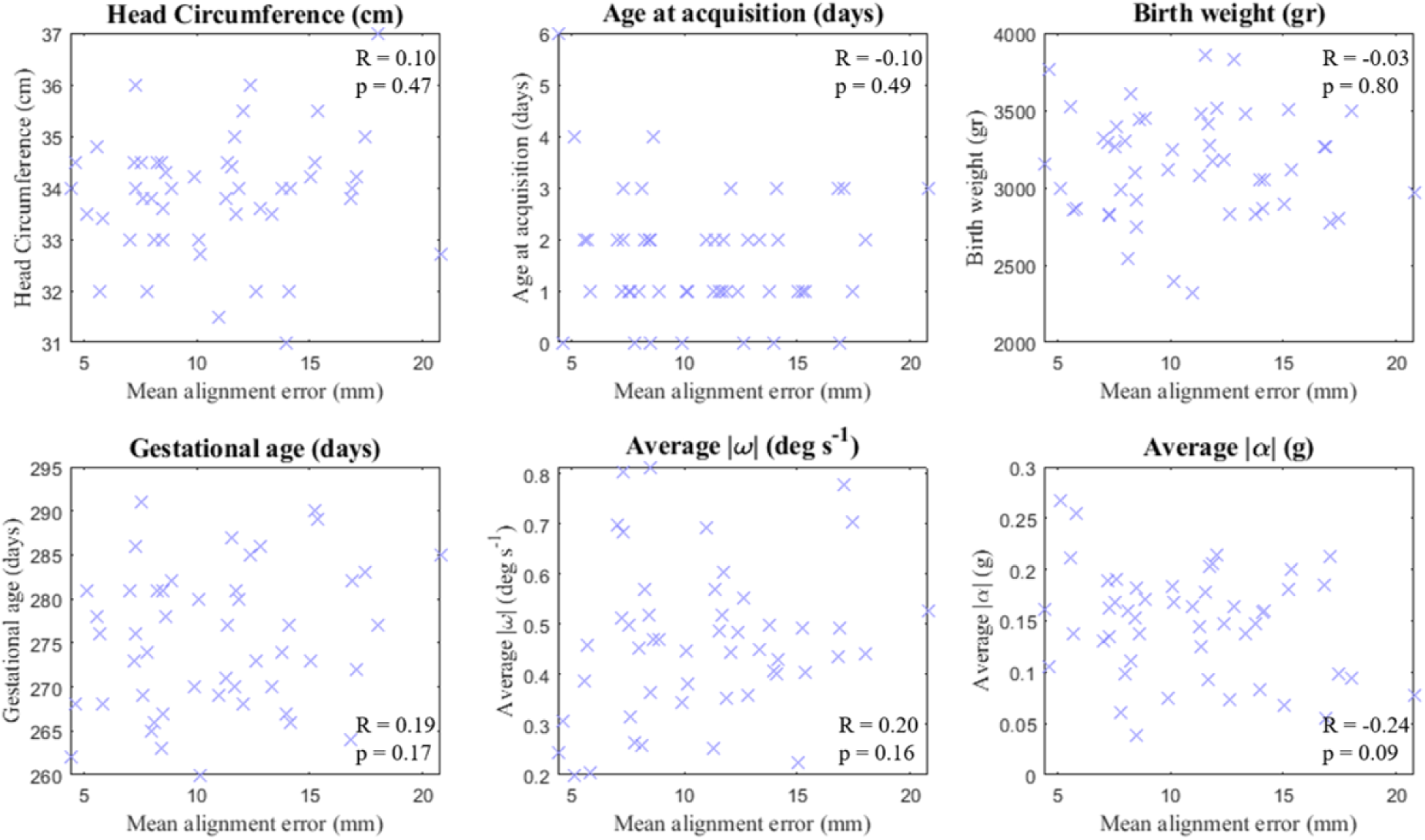
Correlations of newborn metadata and the average probe alignment error between the template and registered probe locations for all subjects. *|ω|* is the magnitude of the angular velocity vector and *|α|* is the magnitude of the angular acceleration vector (units of g, the acceleration due to gravity). No correlations were statistically significant.

## 4 Discussion

Assessing the cognitive functions of newborns in a hospital setting during the first few days of life presents several practical challenges. In particular, testing sessions are often conducted under considerable time pressure, as procedures must remain brief and comfortable for both the infant and the caregiver, they should minimize disruption of clinical care, and avoid prolonged separation from the mother.

A further challenge arises from the newborns’ behavioral state and the associated movement. During testing sessions, infants could transition between different sleep–wake states. Active sleep, in particular, is characterized by frequent body, head, and facial movements. These natural and unavoidable challenges increase the likelihood of cap misalignment in the initial application of the cap and in cap displacement during the experiment and can therefore compromise the consistency of data acquisition procedures. Given the difficulty of collecting neuroimaging data in newborn populations, it is important to develop methods that can account for or correct such imprecisions, rather than discarding otherwise valuable datasets.

In this study, we applied the STORM-NET registration protocol to characterize variability in cap placement and probe displacement across a cohort of 49 newborns, examined its impact on HD-DOT image reconstruction, and evaluated if these displacements were correlated with subject metadata. We reported three representative case studies, the worst case scenario of largest probe displacement, the median case scenario of the median probe displacement and the best case scenario of smallest probe displacement. This approach was intended to provide an interpretable illustration of the range of cap-placement variability observed within the cohort. It also allowed us to examine in detail, how different degrees of probe displacement influence HD-DOT image reconstruction under realistic acquisition conditions while keeping the analysis computationally and visually manageable.

Relative to a centrally aligned template, the cohort exhibited a median probe-position displacement of 10.12 mm which is comparable to the order of magnitude of the image resolution of HD-DOT,^14,15^ demonstrating the importance of accounting for discrepancies between the assumed and reconstructed probe positions. Moreover, by definition, half of the probe displacements are larger than this median value (figure 3) which can become larger compared to the inherit resolution of the HD-DOT, meaning errors in the image reconstruction are compounded by these two factors which affects the localization and spatial accuracy of reconstructed functional responses.

The reason why displacement between a template and subject registered set of probe coordinates causes an error in the reconstructed image, is due to the fact that the forward model which generates in the sensitivity matrix (Jacobian or ‘A’ matrix) requires source-detector locations on a 3D model of the biological tissue being sampled. Since the sensitivity matrix is then used in the image reconstruction and contains all of the spatial information, any deviation from the ground truth of where the source and detector probes cause direct spatial deviations in where functional chromophore changes are reconstructed.

These spatial deviations in chromophore reconstruction are seen most acutely in the most misaligned subject, probe position deviations ranged from approximately 5 mm to 32 mm, with a median of 21.42 mm. These shifts represent meaningful displacements when considered relative to both the physical dimensions of the HD-DOT array and the small size of the newborn head. The adjacent straight line source to detector separation on the Lumo tile used for the HD-DOT in this study are 10 mm 20 mm for the shorter and longer measurement respectively, meaning the largest deviations are comparable to and even exceed the width these probe separations. In addition, on the most misaligned subject, their head circumference was 32.7 cm, meaning the most deviated probe had a deviation of approximately 10% of the head circumference. On an adult size head, the same probe displacement would represent a smaller shift when compared to the subject’s head, highlighting how registration is more important in these developmental populations.

The consequences of this can be seen in figures 6 where probes and tiles are shifted across the surface of the head in the right hand-side direction. The most obvious spatial consequence of this can be observed when considering the two frontal central tiles (left and center tiles respectively in figure 6 B) since in the case of the left tile, all 7 registered probes move from the left hemisphere of the scalp to the right hemisphere or midline compared to the template probe positions, and for the central tile all 7 registered probes are in the right hemisphere of the scalp.

These shifts in probes towards the right hand-side are also observed in the images of changes in HbO, in figure 9, where the clusters and their centroids of HbO reconstructed from the template and registered probe positions are shifted. There are two observations of note to be made here, firstly that the spatial distribution of changes in HbO change in a similar direction due to probe displacement, which can mean interpretation of functional brain responses on the single subject level may be wrong, but more importantly for newborn and infant studies, these spatial discrepancies will propagate up to group-level analyses. Consequently, this may blur or displace functional activation patterns and make results more difficult to interpret.

Secondly, that for the worst case subject, the displacement in probe position is greater than the corresponding displacements in the centroids of HbO. The centroids of positive HbO clusters 1, 3 and 4, which are approximately where the frontal central tile is are shifted by an average of 14.5 mm, whereas the probes in this tile have a displacement of approximately 22 mm. This can be explained since the clusters have spatial properties from the inherent image resolution of HD-DOT and probe displacement discrepancies, the displacement of the centroid of clusters will be and is compounded by these two factors. The spatial resolution will blur the clusters of HbO, which could increase or decrease the distance between adjacent centroids between two clusters of HbO. This will vary across the cortical surface because HD-DOT sensitivity depends on the number and spatial distribution of channels that meet the quality criteria, which in turn are influenced by regional differences in probe–scalp coupling.

In the best and median case subjects, the displacements between the probe positions (figures 4 and 5 respectively) and the centroids of clusters of changes of HbO (figures 7 and 8 respectively) are similar, and smaller than that of the worst case subject. This could be explained by the fact that robe displacements are smaller than the inherent image resolution in these two subjects, meaning the spatial differences in clusters and centroids of HbO are influenced more by image resolution and less the probe displacement, rather than both heavily influenced by image resolution and probe displacement.

Overall, our results show that a registration method such as STORM-NET can help compensate for cap misplacement and resultant in-experiment movement by estimating the actual probe positions for each participant and incorporating them into the HD-DOT pipeline in sensitivity matrix generation. Despite this, since the ground truth positions of probes are not available, it is only possible to compare the registered probe positions with still frames or photographs of the cap. This means that an evaluation of how well the registration was performed can only be done by eye. From the three test case subjects, the general misalignment magnitude and direction was correctly picked up by the registration, it is difficult to truly assess the performance of the registration on a probe-probe level. In general, a more conservative approach of the use of probe registration may be to define acceptable probe displacement thresholds and use these as quality-control criteria for inclusion in image reconstruction or group-level analyses. Participants exceeding these thresholds could either be excluded or analyzed separately.

These findings also raise the question of what factors contribute most strongly to probe displacement and whether they can be systematically controlled. Several factors may contribute to misalignment in newborn HD-DOT studies, including variability in initial cap placement, differences in head size and shape, cap fit, infant movement, behavioral state, and handling during data acquisition. Although we did not observe statistically significant correlations between probeposition error and the variables examined (figure 10), these factors were included because they represent plausible and commonly encountered sources of variability in these populations. In addition, movement variables in average angular velocity and acceleration were included to see if newborns that moved more exhibited cap displacement, but again these correlations were not statistically significant. Better understanding these sources of displacement may help future studies establish more informed quality-control and exclusion criteria aimed at reducing cap misalignment during acquisition.

There are a few limitations of this study, firstly with the focus on the three case study subjects from the entire cohort. The aim of the case studies was not to imply the existence of discrete categories of misalignment, but rather to demonstrate the impact of the full range of probe deviations on HD-DOT. Whilst the probe displacement was calculated on the entire cohort, the more detailed analysis on the affect of this on the HD-DOT was presented using three subjects for two reasons. Firstly, in order to focus the effects of probe displacement on an easy to compare and read number of subjects, as this allowed the study to go into further detail on each of these three subjects in a way that a group level analysis wouldn’t allow for. Secondly, since HD-DOT is computationally intensive and is dependent on individual forward models, performing the HD-DOT twice (template and registered probes) on the entire cohort wouldn’t have offered further insight for the computational cost. Future work could extend this approach to larger subsets of subjects, particularly if subject-specific reconstruction pipelines become necessary for improving spatial accuracy in infant HD-DOT studies.

A further limitation is that optode registration was performed only once, rather than both before and after the recording session. Acquiring registration videos at multiple time points could have provided additional insight into within-subject changes in cap alignment over the course of the experiment. However, implementing this approach in practice proved challenging, as obtaining a clear view of all anatomical and cap reference points required repositioning and lifting the infant’s head, potentially causing additional disturbance and movement. In addition, if performing the registration twice yielded different registered probe positions, then a choice would need to be made to use the first or second set of coordinates, or to use the average of these coordinates.

Finally, there are limitations of the use of the registration technique itself which should be mentioned. The first of these are that the landmark features used in the registration method are only present on the front of the cap, around and between the frontal three tiles, which may cause better registration performance for probes in the front hemisphere of head compared to the rear hemisphere of the head. Whilst the model used for the registration should mitigate this potential problem, if the cap and thus probes are deformed or displaced differently at the rear of the head compared to the front, then less accurate registration may occur in these areas.

## 5 Conclusion

In conclusion, this study demonstrates that cap misalignment is a common source of variability in newborn HD-DOT imaging. The observed probe-position deviations when comparing a centrally aligned template to registered probes were sufficient to change the spatial profile of reconstructed functional images from HD-DOT, highlighting the importance of incorporating subject-specific registration procedures whenever possible. At the same time, the practical challenges inherent to neonatal neuroimaging emphasize the need for acquisition and processing pipelines that are both robust to realistic recording conditions and feasible within sensitive clinical environments. We recommend that future newborn and infant fNIRS and HD-DOT studies use probe registration, report sources of cap misalignment and if they don’t use probe registration, then the steps they perform to ensure reliable and consistent centrally alligned cap placements.

## Code, Data, and Materials Availability

The data for this study can be accessed at https://researchdata.cab.unipd.it/id/eprint/1946 and the code can be found on Github at https://github.com/INMINDteam/HD-DOT-Probe-Registration-Study.

## Acknowledgments

The authors would like to thank the ERC grant IN-MIND 101043216 for funding this project. The first author would also like to acknowledge that we need to break the selective silence on Israel’s genocide in Gaza, as per a recent Lancet correspondence by De Vogli and colleagues urging “professional and academic associations in the fields of health care, public health, and the social sciences to publicly recognize the genocide in Gaza and to revise their official positions accordingly”.^61^ Given this work is in the domain of health care and preterm infants, this statement is entirely relevant here. Genocide by Israel in Gaza was declared by the UN Independent International Commission of Inquiry on the Occupied Palestinian Territory, including East Jerusalem, and Israel (16th September 2025), The International Association of Genocide Scholars (31st August 2025) and by B’Tselem, the Israeli Information Center for Human Rights (27th July 2025). We advocate for peace and prosperity for Israeli’s and Palestinian’s alike.

**Dr Guy A. Perkins** is a postdoctoral researcher at the University of Padova, Italy. His previous work focused on clinical applications using fNIRS and DOT aiming to uncover the neural mechanisms underlying hypo and hyper glycemia in preterm infants. He currently works in the IN-MIND lab which uses HD-DOT for developmental cognitive neuroscience by investigating neural mechanisms of newborn babies verbal memory. Before joining the University of Padova, Dr. Perkins completed his PhD in Physical Sciences for Health at the University of Birmingham, UK, where he specialized in the use of fNIRS and DOT for the assessment of traumatic brain injury and developing multi-frequency frequency domain methods.

**Prof. Silvia Benavides-Varela** is a neuroscientist and an Associate Professor affiliated with the Department of Developmental and Social Psychology and the Padova Neuroscience Center at the University of Padua in Italy. Prof. Benavides-Varela leads the IN-MIND lab and her work bridges, cognitive psychology, and developmental neuroscience, focusing primarily on infant memory, language acquisition, and the neural substrates of mathematical cognition.

Biographies of the other authors are not available.

## List of Figures

1. The source-detector probe set up for the experiment. A. The 45 sources and 60 detectors shown in 2D. These were derived from the raw .lufr file from the LUMO system. B-F. Newborn smoothed gray matter surface model with the source-detector probes and selected EEG (10-5) landmarks shown for reference. These are AFz, AF3, AF4, F7, F3, Fz, F4, F8, T7, C3, Cz, C4, T8, P3, Pz, P4, P7 and P8. B. The right hemisphere. C. Posterior of the cortex. D. Anterior of the cortex. E. The left hemisphere. F. Top view of the cortex.

2. A. The 3D model created and used as part of the Storm-net registration process. B-D. Three frames from the 10 second video required for the Storm-net registration method. These have been redacted as per the requirements for this pre-print.

3. Distributions of the displacement between the template and registered probes (for each given source and detector) for the best, median and worst case scenario subjects respectively, as well as for all 49 subjects. *** denotes a p value of *<* 0.001 from a paired t-test, after corrections for multiple comparisons.

4. The best case subject. A. An image of the best case subject showing the cap alignment, along with lines depicting the anatomically centrally aligned ‘template’ axis from the center of the nose through the central green dot marker and a ‘registered’ axis from the center of the nose through the center of the central frontal tile on the lumo cap. These have been redacted as per the requirements for this pre-print. B-D. The source-detector probes of from the template and from after registration, along with lines showing the displacement between each corresponding source-source and detector-detector, displayed on the surface of the scalp (hidden) above a smoothed gray matter surface head model^58^ (37 week GA).

5. The median case subject. A. An image of the best case subject showing the cap alignment, along with lines depicting the anatomically centrally aligned ‘template’ axis from the center of the nose through the central green dot marker and a ‘registered’ axis from the center of the nose through the center of the central frontal tile on the lumo cap. These have been redacted as per the requirements for this pre-print. B-D. The source-detector probes of from the template and from after registration, along with lines showing the displacement between each corresponding source-source and detector-detector, displayed on the surface of the scalp (hidden) above a smoothed gray matter surface head model^58^ (38 week GA).

6. The worst case subject. A. An image of the best case subject showing the cap alignment, along with lines depicting the anatomically centrally aligned ‘template’ axis from the center of the nose through the central green dot marker and a ‘registered’ axis from the center of the nose through the center of the central frontal tile on the lumo cap. These have been redacted as per the requirements for this pre-print. B-D. The source-detector probes of from the template and from after registration, along with lines showing the displacement between each corresponding source-source and detector-detector, displayed on the surface of the scalp (hidden) above a smoothed gray matter surface head model^58^ (41 week GA).

7. Image reconstruction of ΔHbO from HD-DOT measurements on the best case subject at t = 8 s during a block of auditory stimulation. Each panel A-D shows the location of the significant clusters calculated from using the template probe positions for the Jacobian generation in the first image (highlighted in cyan) and from using the registered probe positions in the second image (highlighted in magenta). The third image in each panel shows the template ΔHbO map overlaid with labels indicating the clusters identified in both the template and registered cases, which are linked with a line for comparative purposes. Labels in pink indicate clusters with significant increments in HbO. Labels in purple indicate clusters with significant decrements in HbO. Each panel shows a different view of the smoothed gray matter surface model, A. Top, B. Posterior, C. Lateral Left and D. Lateral Right. Metrics of these clusters are shown in table 3.

8. Image reconstruction of ΔHbO from HD-DOT measurements on the median case subject at t = 8 s during a block of auditory stimulation. Each panel A-D shows the location of the significant clusters calculated from using the template probe positions for the Jacobian generation in the first image (highlighted in cyan) and from using the registered probe positions in the second image (highlighted in magenta). The third image in each panel shows the template ΔHbO map overlaid with labels indicating the clusters identified in both the template and registered cases, which are linked with a line for comparative purposes. Labels in pink indicate clusters with significant increments in HbO. Labels in purple indicate clusters with significant decrements in HbO. Each panel shows a different view of the smoothed gray matter surface model, A. Top, B. Posterior, C. Lateral Left and D. Lateral Right. Metrics of these clusters are shown in table 4.

9. Image reconstruction of ΔHbO from HD-DOT measurements on the worst case subject at t = 8 s during a block of auditory stimulation. Each panel A-D shows the location of the significant clusters calculated from using the template probe positions for the Jacobian generation in the first image (highlighted in cyan) and from using the registered probe positions in the second image (highlighted in magenta). The third image in each panel shows the template ΔHbO map overlaid with labels indicating the clusters identified in both the template and registered cases, which are linked with a line for comparative purposes. Labels in pink indicate clusters with significant increments in HbO. Labels in purple indicate clusters with significant decrements in HbO. Each panel shows a different view of the smoothed gray matter surface model, A. Top, B. Posterior, C. Lateral Left and D. Lateral Right. Metrics of these clusters are shown in table 5.

10. Correlations of newborn metadata and the average probe alignment error between the template and registered probe locations for all subjects. *|ω|* is the magnitude of the angular velocity vector and *|α|* is the magnitude of the angular acceleration vector (units of g, the acceleration due to gravity). No correlations were statistically significant.

## List of Tables

1. A summary of the demographic and clinical information, for the full sample, the best case subject, median case subject and worst case subject in terms of probe alignment respectively. Gestational age (GA) given by the number of weeks (W) plus the number of days (D). BW = Birth weight in grams. Head Circ. = Head Circumference in centimeters. Values presented are in the format median (IQR 1-3), where appropriate.

2. The optical properties used for the different tissues in the mesh Finite Element Model (FEM).^16^ CSF = cerebral spinal fluid, GM = gray matter, WM = white matter. All values given in units of mm*^−^*^1^.

3. Metrics for each cluster of ΔHbO from the best case subject in figure 7. These metrics are the displacement between the centroid of the ΔHbO clusters from the template to the registered images (euclidean distance) and the dice coefficient between these.

4. Metrics for each cluster of ΔHbO from the median case subject in figure 8. These metrics are the displacement between the centroid of the ΔHbO clusters from the template to the registered images (euclidean distance) and the dice coefficient between these.

5. Metrics for each cluster of ΔHbO from the best case subject in figure 9. These metrics are the displacement between the centroid of the ΔHbO clusters from the template to the registered images (euclidean distance) and the dice coefficient between these.

